# “Comparative Analysis of Ovarian Transcriptome Changes Across Gestational Stages in Kari Sheep”

**DOI:** 10.1101/2023.11.20.567795

**Authors:** Haidar Ali, Sohail Ahmad

**Author notes:** Corresponding Author: Haidar Ali, Institute of Biotechnology and Genetic Engineering, The University of Agriculture Peshawar 25000, Pakistan.

## Abstract

This study aimed to investigate the genetic determinants of gestation length in Kari sheep, employing RNA-Seq technology. Employing a comprehensive whole transcriptome analysis, we sought to pinpoint differentially expressed genes (DEGs) while also delving into gene ontology (GO) enrichment and Kyoto Encyclopedia of Genes and Genomes (KEGG) pathway assessments. The analysis revealed the identification of a total of 19,546 genes expressed in ovary. While comparing the transcriptomes of Kari sheep with Balkhi, yielding 976 DEGs (p < 0.05, Log2fc>1, <-1). Notably, among these DEGs, an upregulation of genes was observed associated with Ubiquitin-protein transferase activity, such as *CNOT4, RC3H1*, and *XIAP*. Concurrently, DEGs like *NFAT5, EPAS1, ZNF644, RBPJ*, and *FOXP2* exhibited associations with RNA polymerase II core promoter proximal region sequence-specific DNA binding. Conversely, downregulated genes, including *EEA1, CNOT4, FGD4, MBNL1, ZRANB2, REV3L, XIAP, ATP13A3, RPAP2, FOXP2*, and ADAMTS6, were implicated in the mRNA surveillance pathway. In addition, several Gene Ontology terms, such as GO:0001228 (transcriptional activator activity) and GO:0004842, along with GO:0000978 (transcriptional activator activity), were linked to the DEGs. KEGG pathways, including “Glycosaminoglycan biosynthesis - chondroitin sulfate/dermatan sulfate” (KEGG:532) and “basal cell carcinoma” (KEGG:5217), were associated with our findings. Our principal component analysis (PCA) demonstrated a cohesive clustering of gene expression profiles among the four samples, with subtle distinctions. Protein-protein interaction (PPI) analysis indicated the functional relationships among the DEGs. Notably, genes such as *ABHD16B* and *NPBWR2* exhibited strong co-expression among the down-regulated DEGs, while *DNAH7/TBC1D31* and *MBNL1/NOVA1* displayed prominent co-expression among the up-regulated DEGs. Consequently, our study offers a comprehensive understanding of Kari sheep genetics and the pivotal genes involved in gestation length determinants. These findings carry significant genetic implications, enhancing genetic resources, furthering reproductive biology comprehension, and contributing to the advancement of sustainable sheep farming practices.

## INTRODUCTION

Livestock plays a pivotal role in Pakistan’s burgeoning economy, with sheep emerging as a significant and abundant domesticated species. Within the diverse spectrum of sheep breeds in Khyber Pakhtunkhwa (KP), a region of rich biological diversity, the Kari sheep breed stands out as a noteworthy subject of study. This breed, which encompasses distinct characteristics, holds a unique place in the local landscape.

Kari sheep, a small-sized and thin-tailed breed with a predominant white coat color, was discovered in the remote “Lotkho” area of Chitral, KP, during a survey conducted in 2000. While Kari sheep possesses distinct features that set it apart, the most intriguing characteristic lies in its remarkably short gestation period. This breed’s gestation length, reported to range from 92 to 120 days, differs significantly from the broader ovine population. Such a unique reproductive trait demands comprehensive investigation to unravel the underlying molecular mechanisms.

Reproductive efficiency is a key-determinant character of sheep breeding systems, determined by the lambing interval and litter size further affected by a-seasonality in lambing pattern, gestation length and dry period standing at the forefront. The economic prosperity of sheep farming is intricately linked to the number of lambs weaned and their average weight per ewe per year.

Despite the importance of Kari sheep’s reproductive qualities, reports on variations in gestation length have been relatively scant. Gestation length, a multifaceted trait, is known to be influenced by fetal, maternal, genetic, and environmental factors. A broader understanding of these factors, encompassing genetic, external, and internal environmental aspects, is crucial to decipher the intriguing variations in gestation length observed in Kari sheep.

In this context, gestation length is one of the biological fix period for the entire specie, however in the current study it emerges as another most important reproductive trait variable, profoundly influenced by genetic and environmental factors. The succinct gestation period of Kari sheep sparks curiosity and warrants a thorough exploration of the genetic underpinnings contributing to this distinct attribute. In the current study, the Balkhi sheep breed served as a control for possessing a specie representative (normal) gestational duration (158-160 days) in comparison to Kari.

Traditionally, investigations into gene expression relied on low-throughput techniques, limiting the study of one transcript at a time. However, the past two decades have witnessed the advent of high-throughput sequencing methods, particularly RNA-Seq technology, enabling comprehensive transcriptome profiling. This technological advancement has revolutionized our ability to explore diverse physiological and pathological states by providing a holistic view of gene expression.

In this groundbreaking study, we harnessed high-throughput sequencing to examine into the entire transcriptome of Kari sheep throughout their gestational journey, with a particular focus on identifying differentially expressed genes (DEGs). The primary objective of this research endeavor was to enrich the genetic resources of Kari sheep by capturing the complete gene sequences associated with gestational duration and other pivotal biological processes during ovarian development. Significantly, this study represents the first of its kind, shedding light on the ovarian transcriptomes of Kari sheep. Our findings are poised to serve as a foundational cornerstone for further explorations into functional transcriptomics within this unique breed, contributing to an enhanced understanding of its genetic dynamics and reproductive biology

## MATERIAL AND METHODS

This study was carried out at the Animal Biotechnology Laboratory, Institute of Biotechnology and Genetic Engineering (IBGE), the University of Agriculture Peshawar, KP. The selection of Kari sheep for this study is due to their unique characteristics such as small body length and short gestation period. Moreover, Kari sheep predominantly inhabits the hilly regions of Chitral, characterized by significantly lower temperatures. Conversely, the inclusion of Balkhi sheep with larger body and longer gestation as well Balkhi sheep represent a common domesticated breed in Afghanistan and northwestern Pakistan. This comparative transcriptome study is set to reveal the valuable insights into genetics of genes regulating the short gestation period found in both breeds.

### ANIMAL SELECTION AND BLOOD SAMPLING

Kari sheep was standing as a core breed whereas Balkhi breed served as a control. Animals were synchronized using various hormonal disruptive medications such as Prostenol and Conceptal. Sheep gestation was categorized into three trimesters but sampling was done from 1^st^ and 3^rd^ trimesters. For transcriptomic, a total of ten animals (Biological replicates (BR) n=10: Technical replicates (TR) n=2) from each breed was selected in each trimester. The comparison in the groups [Biopsy Balkhi (BB) vs Biopsy Kari (BK)] were done based on the respective trimester. Biopsy samples were taken through ovariectomy, snap frozen in -80°C until RNA extraction.

**Table 1.**
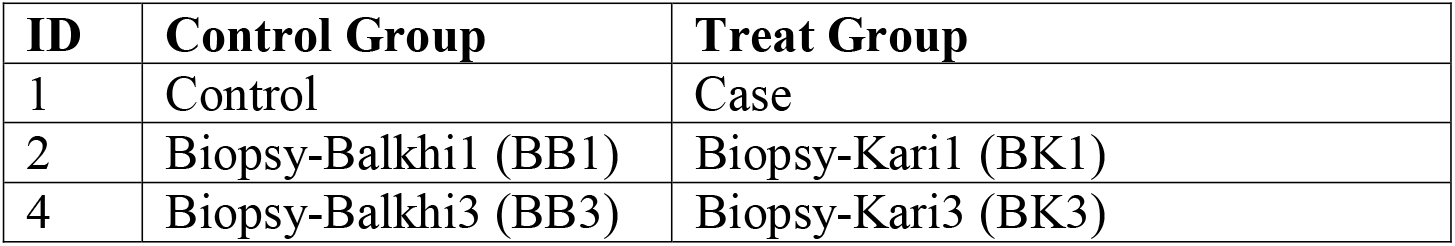
Group of sheep used during transcriptome analysis for gestational length.

### RNA EXTRACTION, LIBRARY CONSTRUCTION, AND SEQUENCING

Total RNA was extracted through TRiZol method [11] and quantified through NANODROP (IMPLEN, CA, United States), OD260/OD28 ratio greater than 1.7 was selected for deep sequencing. RNA was reverse-transcribed into cDNA using RevertAid cDNA synthesis Kit and was sent to BGI (https://www.bgi.com/global) China for high-throughput sequencing. A total of 4 samples were tested using the DNBSEQ platform, with an average yield of 6.75G data per sample (PE150).

### DATA PROCESSING

The acquisition of clean data required the removal of reads including adapters or ploy-N, as well as the removal of low-quality reads from raw data. FastQC software was used for this process. At the same time, Q20, Q30 and GC content of the clean data were calculated. Clean reads were mapped to the reference genome (GCF_000298735.2_Oar_v4.0, https://www.ncbi.nlm.nih.gov/datasets/genome/GCF_016772045.1). After getting clean reads, we used HISAT (https://ccb.jhu.edu/software/hisat2/index.shtml) (Kim et al., 2015) to align the clean reads to the reference genome. Align clean reads to the reference genes by using Bowtie2 (https://bio.tools/bowtie2) to get the alignment result. For samples with good quality and sufficient sequencing data, most of the transcripts were completely covered, and reads were evenly distributed in various regions of the transcript.

### DIFFERENTIAL EXPRESSION ANALYSIS AND FUNCTION ANNOTATION

Analysis of each group for DEGs was performed using DESeq2 R programme. The feature Countssoftware removed counts below 5 in every sample to improve precision. The pair-wise false discovery rate (FDR) of each gene was calculated using the Benjamini-Hochberg method. Differential gene expression cutoffs were p-values less than 0.05 and log2fold changes higher than 1. ShinyGO online database (<u>ShinyGO 0.77 (sdstate.edu)</u>) was used to perform PCA on FPKM values of BB and BK. The (eu-biosys.bgi.com/#/main) online tool was used to study differentially expressed genes’ biological roles. Gene Ontology (GO) functional enrichment analysis for cellular components (CC), biological processes (BP), and molecular functions (MF) as well as KEGG pathway analysis was performed using DAVID online tool (<u>DAVID Functional Annotation Tools</u> (ncifcrf.gov)). The analysis was done on (eu-biosys.bgi.com/#/main).GO and KEGG pathway functional enrichment analysis (Fisher’s exact test) cutoff criteria was p-value of less than 0.05. The expression trends graph was designed using SR-Plot online database (http://www.bioinformatics.com.cn/srplot). PPI were created using String online database (<u>STRING: functional protein association networks</u> (string-db.org)) and Cytoscape (Version 3.10.1). The Pearson Correlation was measured using SPSS (Version 20)

## RESULTS

The average GL of Kari was 100 ± 5 days while that of Balkhi was 147± 5 days. To obtain a global overview of genes related to sheep gestational length, we performed pairwise comparisons BB1 vs. BK1, BB3 vs. BK3. Four separate cDNA libraries were constructed from the biopsy. A total of 187.56 million raw reads were obtained from biopsy, the total number of bases for each of the samples varied from 44.18 to 45.08 Gb. After removing adapters, low-quality and low-complexity reads, high-quality RNA sequencing data were generated. Subsequently, 26.94 Gb clean reads were obtained, with each sample having >6.63 Gb. The Q20, Q30 of the clean data were simultaneously calculated. The Q20 (the percentage of bases with a Phred score greater than 20) and Q30 (the percentage of bases with a Phred score greater than 30) were higher than 94%, and the mean average GC content was 47.8%. The ratio of reads mapping uniquely to the reference sheep genome was ranged from 76.19% to 84.08% (Table.2). All the statistical information has been attached in Supplementary file Table S1_Basic info.

**Table 2.** Statistical information of RNA Seq data in each sample.

| Sample | Total Raw Reads (M) | Total Clean Reads (M) | Total Clean Bases (Gb) | Clean Reads Q20 (%) | Clean Reads Q30 (%) | Clean Reads Ratio (%) | GC % | Uniquely Mapping (%) |
| --- | --- | --- | --- | --- | --- | --- | --- | --- |
| BB1 | 45.57 | 44.18 | 6.63 | 98.16 | 94.41 | 96.95 | 51.1 | 76.19 |
| BK1 | 47.33 | 44.88 | 6.73 | 97.94 | 94 | 94.82 | 43.4 | 78.48 |
| BB3 | 47.3<br>3 | 45.44 | 6.82 | 98.15 | 94.37 | 96.01 | 50.<br>8 | 85.91 |
| BK3 | 47.3<br>3 | 45.08 | 6.76 | 98.04 | 94.02 | 95.25 | 45.<br>9 | 84.08 |
| Total | 187.<br>56 | 179.58 | 26.94 | 392.29 | 376.8 | 383.03 | 47.<br>8 | 324.66 |

### DIFFERENTIALLY EXPRESSED GENES ANALYSIS

A total of 19546 genes were found in the biopsy of each sample. DESeq2 R package was used to obtain DEGs. A total of 976 DEGs were found when comparing transcriptomes of Kari and Balkhi sheep (p < 0.05) Log2fc>1 (Supplementary File_Table S2). Of them, 70 genes were upregulated (URGs) and 476 genes were downregulated (DRGs) in BK1 vs. BB1. In BK3 vs. BB3, the number of upregulated genes were 6 while the downregulated were 177 respectively. Additionally, 113 genes were commonly expressed in all the samples.

### GO ENRICHMENT ANALYSIS OF DEGs

GO enrichment analysis was performed to shed light on the potential function of DEGs concerned with gestational length in both groups. In the study, we separated the up- and down-regulated genes for GO analysis and found that in BK1vs.BB1, many Up-regulated DEGs enriched GO terms related to biological processes: Positive regulation of cardiac muscle cell proliferation (GO:0060045), Blood vessel remodeling (GO:0001974), Synapse assembly (GO:0007416), and Ubiquitin-dependent protein catabolic process (GO:0006511). And some up-regulated genes were associated with cellular components, such as Integral component of presynaptic membrane (GO:0099056), Nucleoplasm (GO:0005654), Cytosol (GO:0005829), Cytoplasmic vesicle (GO:0031410), and Nucleolus (GO:0005730). Other up-regulated genes were associated with molecular functions such as Transcriptional activator activity (GO:0001228), Ubiquitin-protein transferase activity (GO:0004842), RNA binding (GO:0003723), RNA polymerase II core promoter proximal region sequence-specific DNA binding (GO:0000978) and Metal ion binding (GO:0046872). Similarly, in BK1vs.BB1, many Down-regulated DEGs enriched GO terms related to biological processes: Protein ubiquitination (GO:0016567), Behavioral fear response (GO:0001662), Negative regulation of cell growth (GO:0030308), Negative regulation of signal transduction (GO:0009968) and Visual learning (GO:0008542). And, some down-regulated genes were associated cellular components such as: Voltage-gated potassium channel complex (GO:0008076), Extracellular exosome (GO:0070062), Cell junction (GO:0030054), Integral component of plasma membrane (GO:0005887) and Perinuclear region of cytoplasm (GO:0048471). Other down-regulated genes were associated with molecular functions: Ubiquitin protein ligase binding (GO:0031625), Guanyl-nucleotide exchange factor activity (GO:0005085), Heparin binding (GO:0008201), Beta-galactoside (CMP) alpha-2,3-sialyltransferase activity (GO:0003836) and Neuregulin binding (GO:0038132). In BK3vs.BB3, we found many Down-regulated DEGs enriched GO terms related to biological processes: Osteoblast development (GO:0002076), Endocytosis (GO:0006897), Negative regulation of endothelial cell migration (GO:0010596), Negative regulation of angiogenesis (GO:0016525) and Peripheral nervous system development (GO:0007422). And some down-regulated genes were associated with Cellular functions such as Axon initial segment (GO:0043194), Schaffer collateral - CA1 synapse (GO:0098685), Intercellular bridge (GO:0045171), Cell surface (GO:0009986) and nuclear body (GO:0016604). Other down-regulated genes were associated with molecular functions: GTPase activator activity (GO:0005096) and Phosphatidylinositol binding (GO:0035091). The top 05 Go terms for BK1vsBB1 and BK3vsBB3 are shown in (Figures 2 A-F). In BK3vs.BB3, the up-regulated genes were only six in numbers. Their role in biological processes, cellular components and molecular function are given in the below Table3. Statistical results of all GO terms are given in Supplementary file (Table S3-S6).

**Figure 1.**
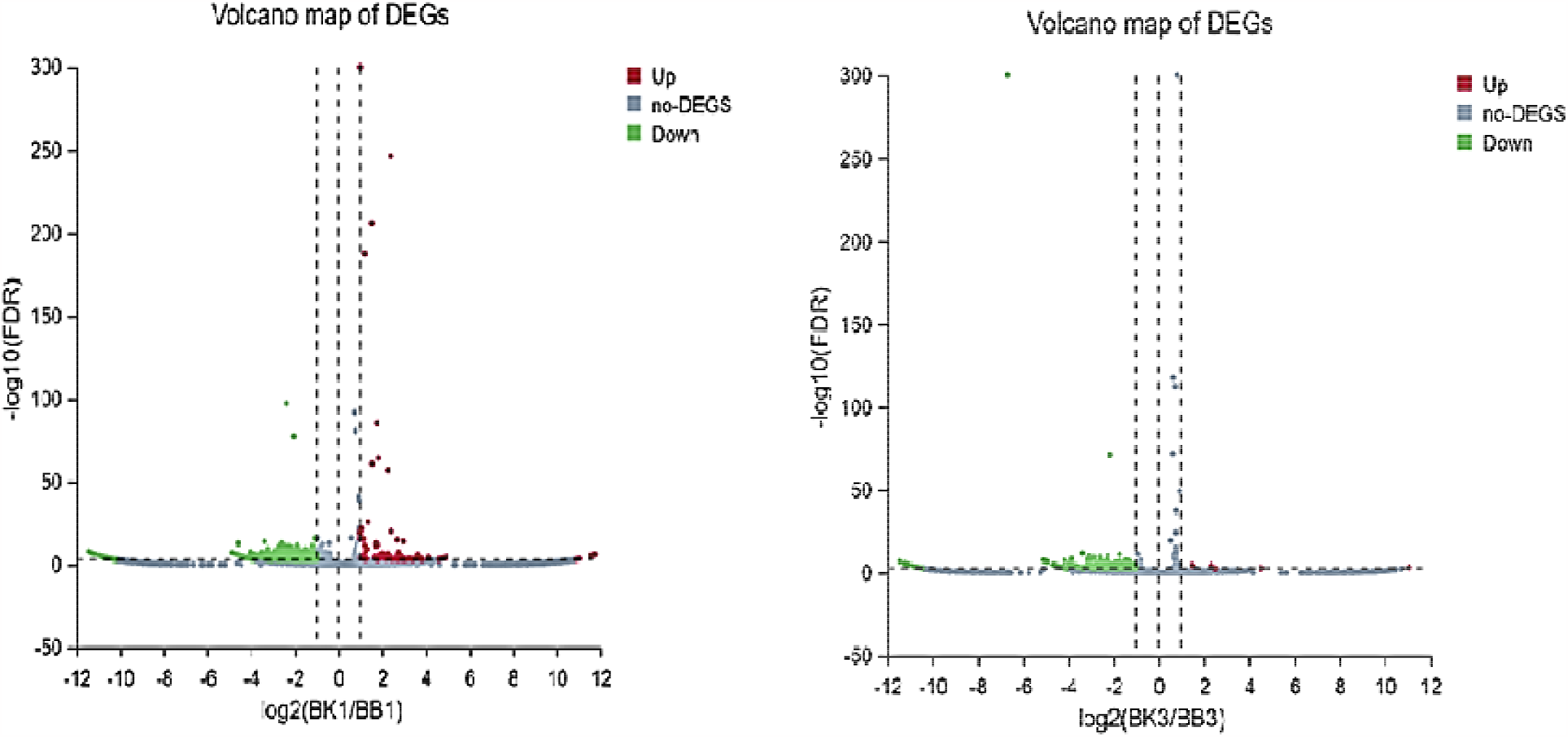
Volcano Plot of Up and Down-regulated genes in two breeds. The vertical represents the fold change (BK1vs.BB1, BK3vs.BB3) ≥1 and ≤ respectively. And the horizontal axis shows the adjusted p-value (FDR q-value) of 0.05. The color dots represent the FDR (q-value). A) DEGs in BK1vs.BB1; B) DEGs in BK3vs.BB3.

**Figure 2.**
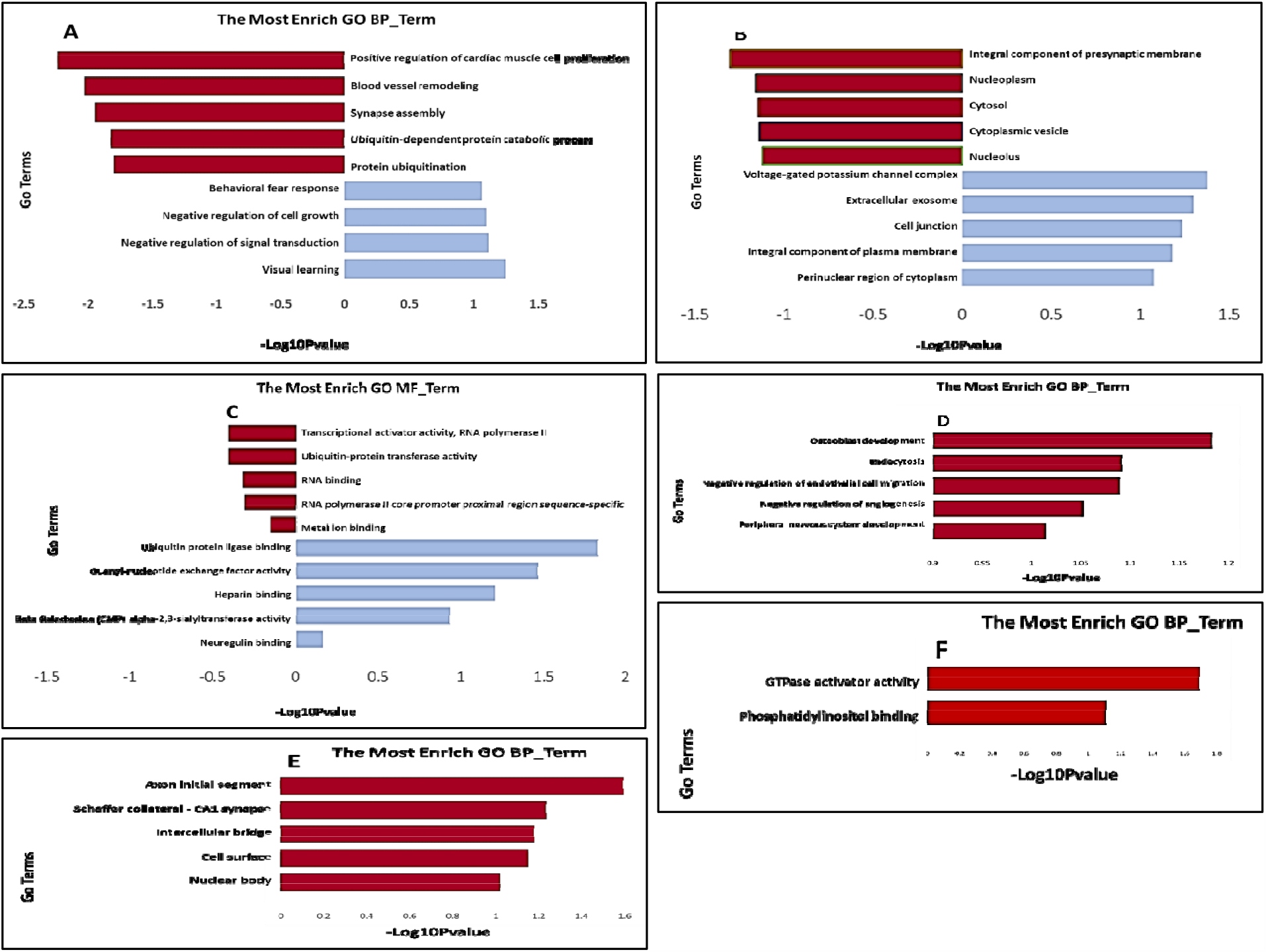
Significantly enriched Gene Ontology (GO) terms of DEGs based on their functions. The Bar Graph of top 05 BP (A), CC (B) and MF (C) terms in the enrichment analysis of DEGs in the BK1vsBB1 URGs. Bar Graph of the top 05 BP (A), CC (B) and MF (C) terms in the enrichment analysis of DEGs in the BK1vsBB1 DRGs. Bar Graph of the top 05 BP (D), CC (E) and MF (F) terms in the enrichment analysis of DEGs in the BK3vsBB3 DRGs. BP: biological process; CC: cellular component; MF: molecular function. “Red” represents for GO terms of down-regulated genes and “Blue” represents for GO terms of up-regulated genes in DP.

**Table 3:**
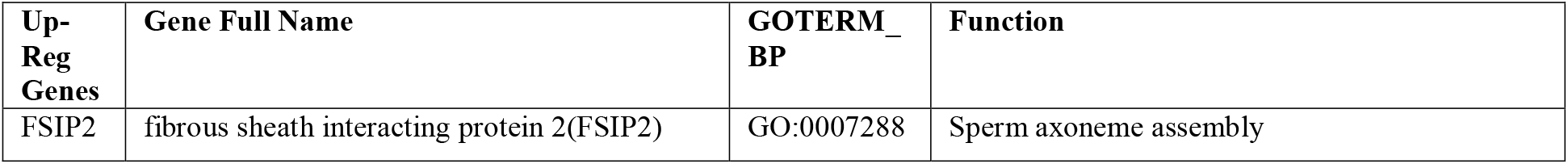
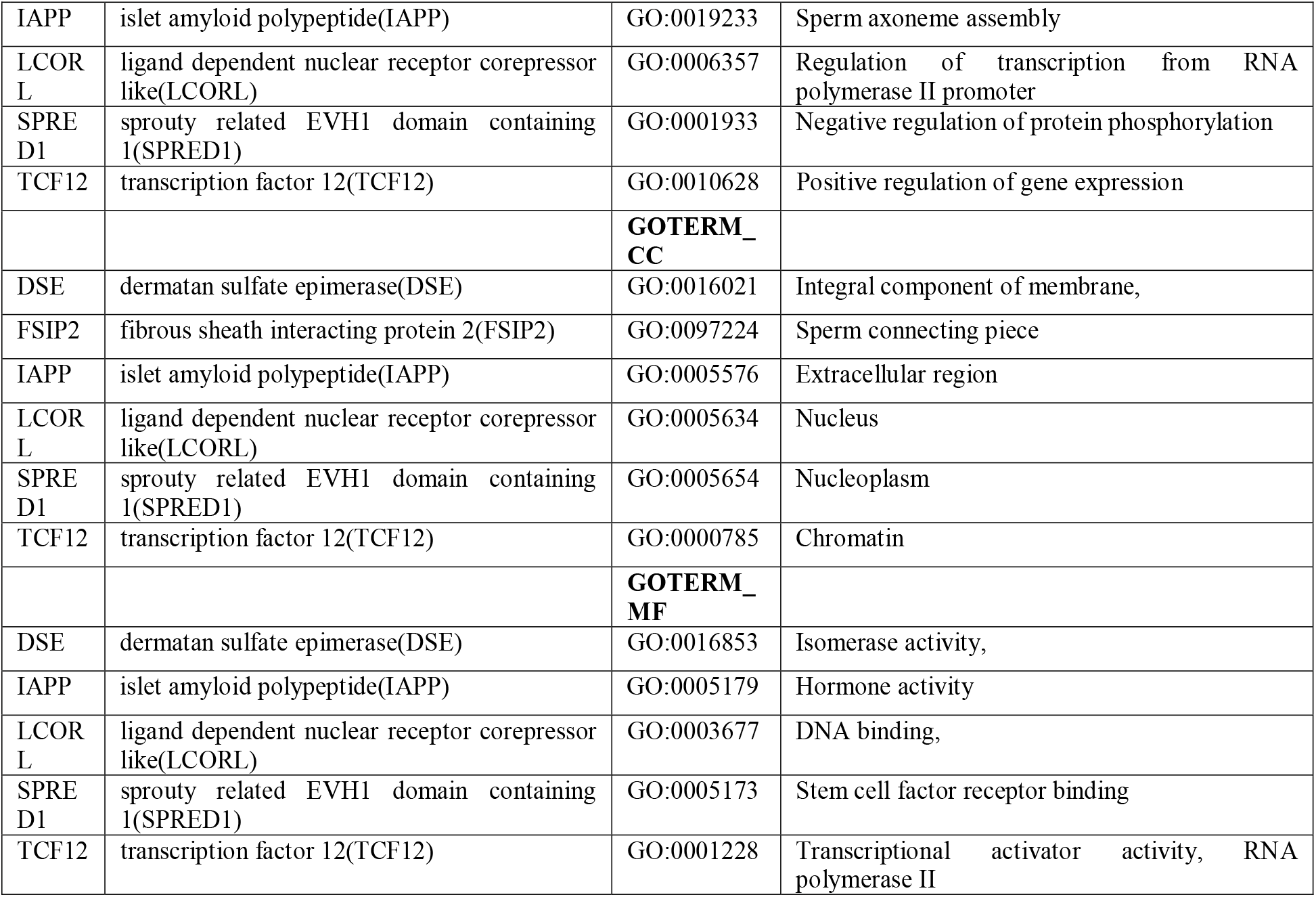
Upregulated genes, Go terms and functions of DEGs in the BK3vsBB3.

### KEGG PATHWAYS ANALYSIS OF DEGs

Based on the KEGG pathway database, we could systematically analyze gene function and molecular networks in cells. In the current study, those 477 DEGs were assigned to KEGG pathways. Out of 234 KEGG pathways in the BK1vsBB1, down-regulated DEGs were annotated to Metabolism Glycosaminoglycan biosynthesis - chondroitin sulfate / dermatan sulfate (532), Cancer: specific types including Basal cell carcinoma (5217), Sensory system such as Phototransduction (4744), Cellular community - eukaryotes Signaling pathways regulating pluripotency of stem cells (4550), and Endocrine system including GnRH secretion (4929), etc. The up-regulated DEGs of the Bk1vsBB1 were mainly enriched in several metabolic pathways including mRNA surveillance pathway (3015), D-Amino acid metabolism (470) and Inositol phosphate metabolism (562). In addition, Drug resistance: antineoplastic such Platinum drug resistance pathway (1524) and Phosphatidylinositol signaling system (4070) were also significantly enriched.

A total of 174 KEGG pathways were annotated in the BK3vsBB3, and the down-regulated genes were mainly associated with important signaling pathways and organismal systems such as Cushing syndrome (4934), Glycan biosynthesis and metabolism (533), Endocrine system (4929) and Cortisol synthesis and secretion (4927). Meanwhile, up-regulated genes are associated with Glycan biosynthesis and metabolism (532), Endocrine and metabolic disease (4950) and Signaling molecules and interaction (4080). The top 40 significant pathways (20 for down-regulation, 20 for up-regulation) in each group were shown in (Figures 3 A-D), and all the significant KEGG results were shown in Supplementary File. Statistics for KEGG results for each group were available in Supplementary file (Table S7-S10).

**Figure 3:**
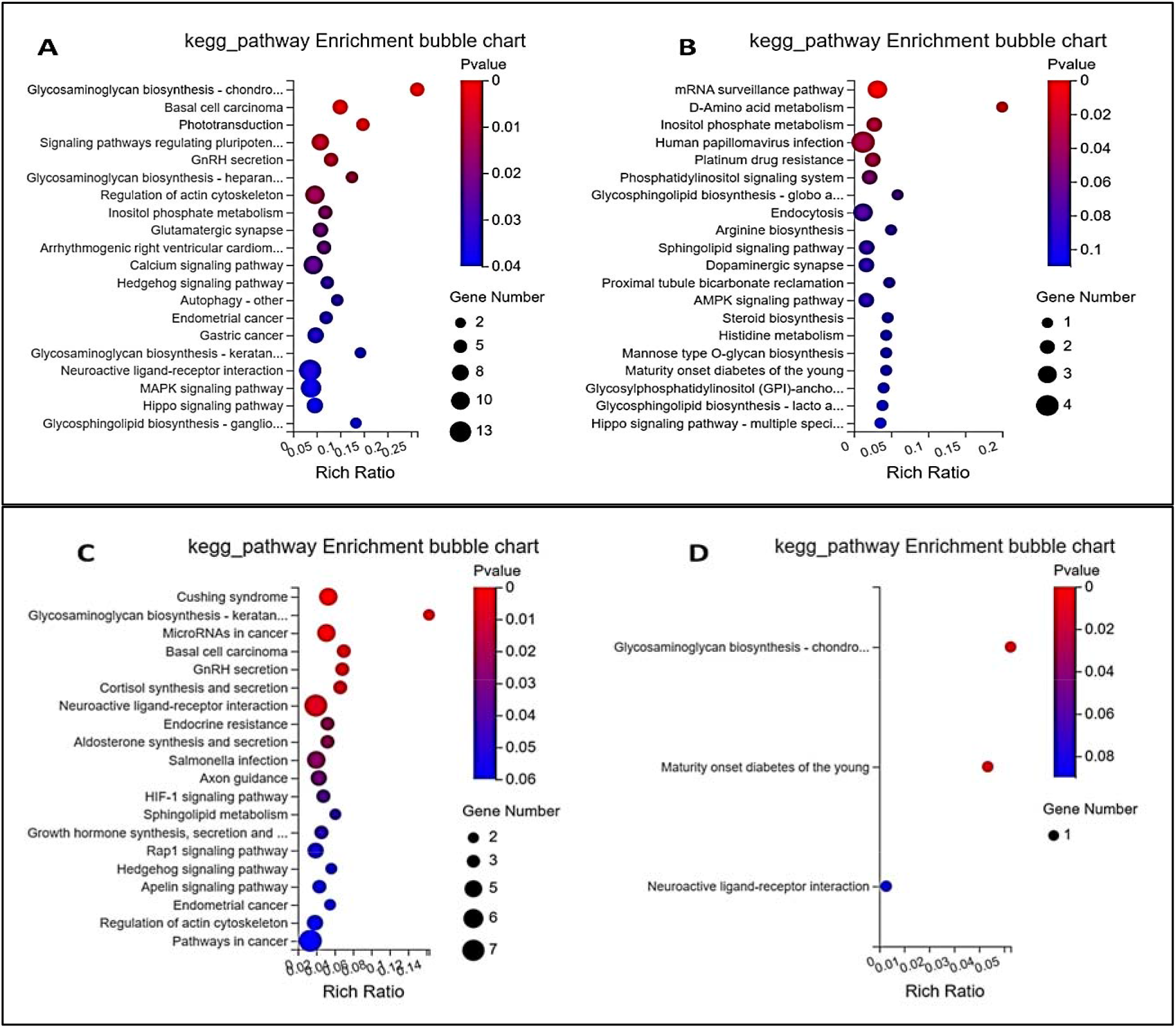
Bubble Plots showing the functional pathways of KEGG enrichment: The top 20 pathways shown in (A) pathways of down-regulated genes of BK1vsBB1; (B) pathways of up-regulated genes of BK1vsBB1; (C) pathways of down-regulated genes of BK3vsBB3 and (D) pathways of up-regulated genes of BK3vsBB3.Up-and Down-regulation refers to gene expression in BKvsBB.

### SCATTERED PLOT SHOWING DEGS EXPRESSION

The average expression of DEGs in each group is shown in figure 4 (A-E). The R value in a scatter plot represents the correlation coefficient between two variables. It is a measure of the strength and direction of the linear relationship between two variables. The R value between BB1 and BK1 was (R=0.06) while between the BK1vsBK3, the R-value was (R=0.86). The R value between BB1 and BB3 was (R=0.89) while between the B3 and BK3, the R value wa (0.74). The average DEGs expression in each group was shown in figure 4E, where the highly Up-and Down-regulated genes are nominated with arrows. All the statistical data have been attached (Supplementary File_Table S11-12)

**Figure 4:**
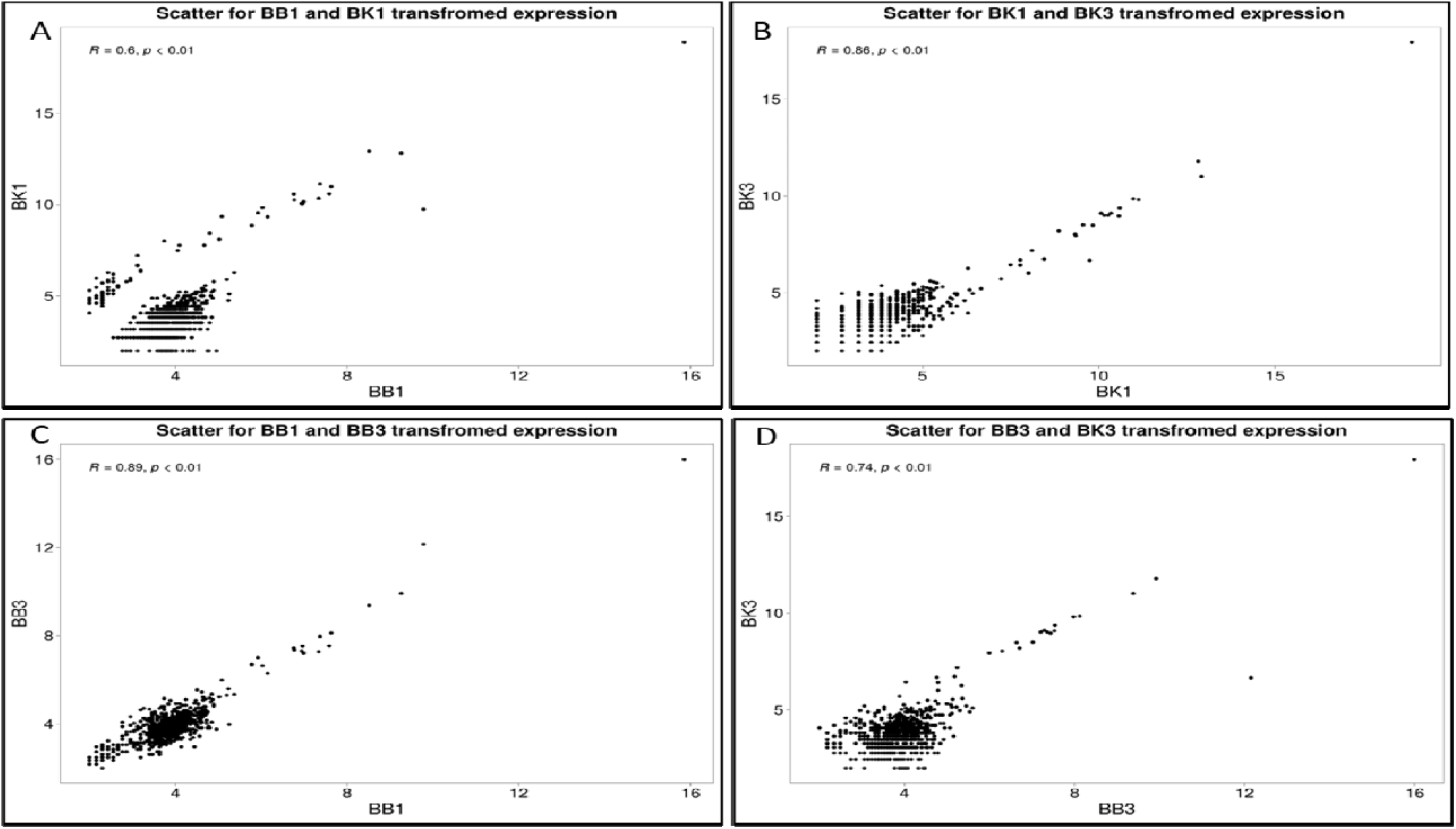
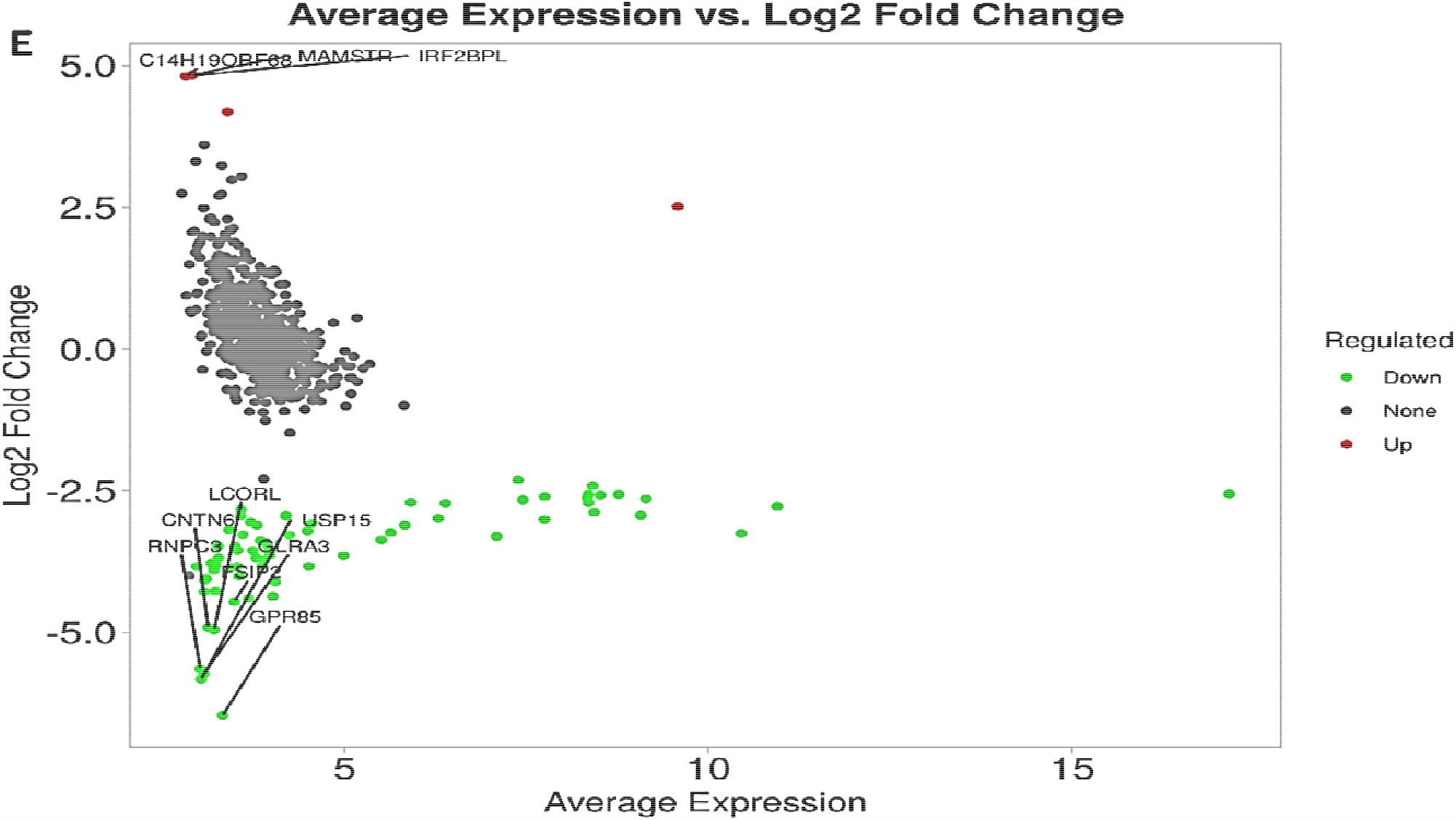
Showing the scattered Plot of DEGs expressions in each group. (A) represents the average expression between BB1vs.BK1, (B) represents the average expression between BK1vs.BK3, (C) the average expression between BB1vsBB3, (D) represents the average expression between BB3vs.BK3, and (E) shows the average expression of DEGs all groups. Highly Up-and Down-regulated genes are mentioned in Figure4E.

### BOXPLOT EXPRESSION QUANTITY OF DEGS

In this analysis, we compare two gene sets: Downregulated Genes (DRGs) and Upregulated Genes (URGs), represented by boxplots “A” and “B” respectively. Boxplot “A” shows that DRGs had higher expression levels in the control group than in the experimental condition, indicating reduced expression in response to the experiment. Conversely, boxplot “B” demonstrates that URGs had higher expression in the experimental group, indicating increased expression under specific conditions. The statistical data for boxplot have been attached (Supplementary File_Table S13)

**Figure 5:**
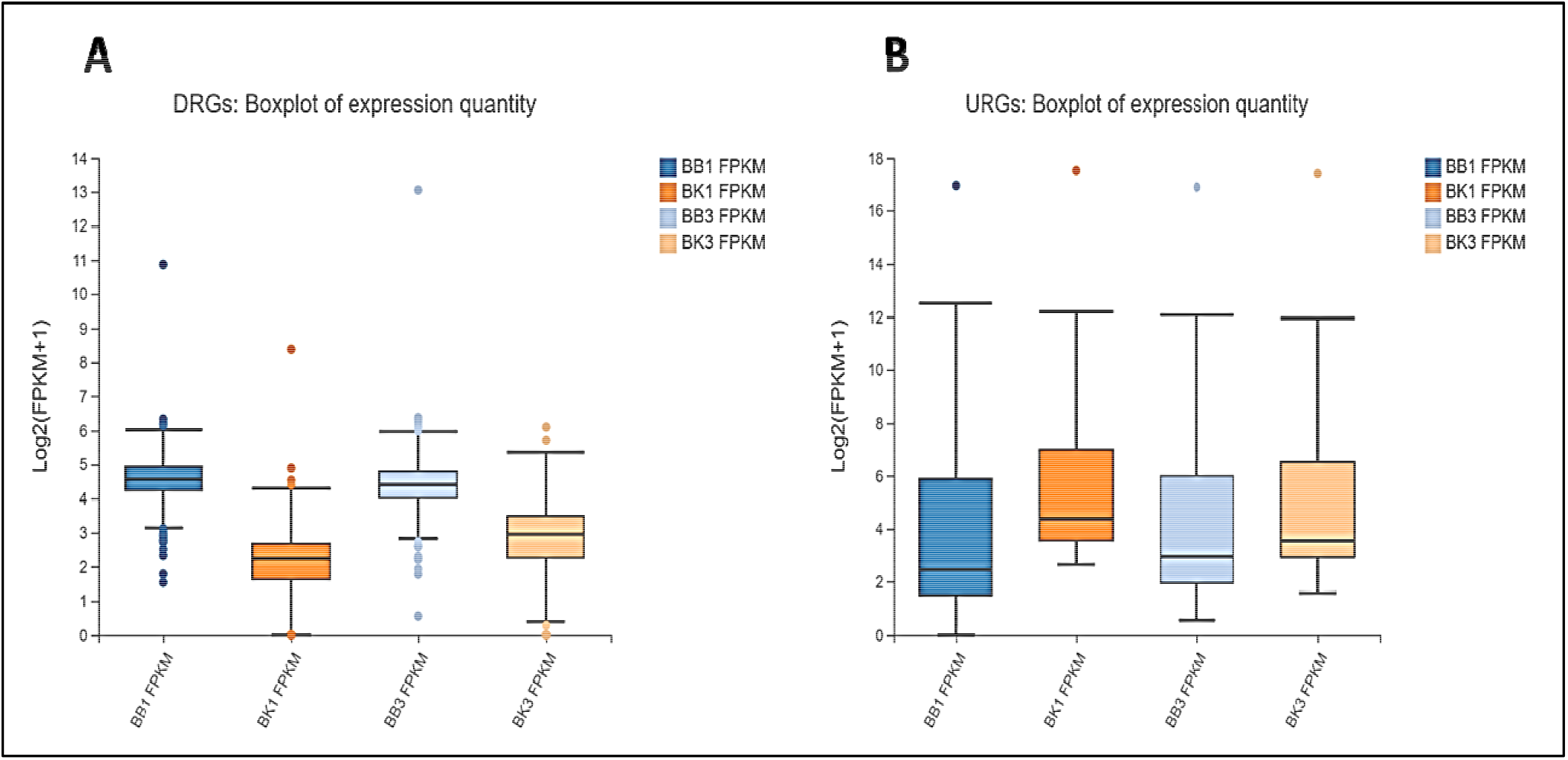
boxplots showing a visual comparison, highlighting the differential gene expression between DRGs (A) and URGs (B) in response to the experimental conditions in BB vs BK.

### COEXPRESSION TREND BETWEEN DEGs_BBvs.BK

In examining gene expression trends for four samples (BK1, BB1, BB3, BK3), different patterns emerge for up-regulated and down-regulated genes. This pattern reveals complex changes in gene function during pregnancy in Kari and Balkhi sheep, and reveals gene regulation during pregnancy This study provides valuable insights value in the molecular mechanisms associated with gestation in these two breeds of sheep.

**Figure 6:**
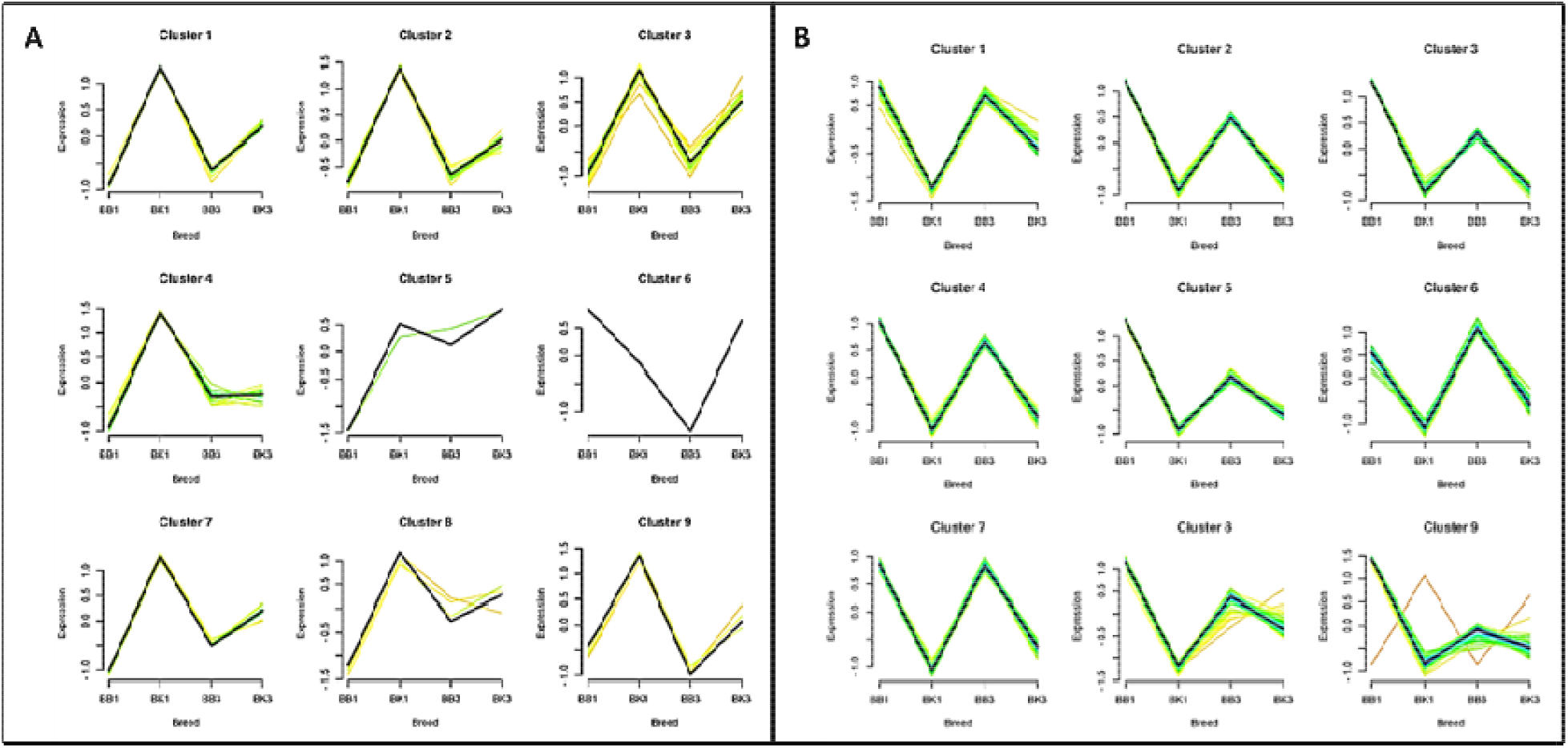
showing the gene expression trends for four samples (BK1, BB1, BB3, BK3), different patterns emerge for up-regulated (A) and down-regulated (B) genes.

### PCA ANALYSIS AMONG BB AND BK

PCA analysis visually identifies classifications and relationships based on gene expression of two breeds of sheep BB (Balkhi) and BK (Kari) Each point in the PCA plot represents a model, the construction of these points there reflects the degree of similarity or similarity between the two groups. the PCA Component 1 values for all four samples (BB1, BB3, BK1, and BK3) are very close, indicating that they have similar gene expression profiles along the first principal component axis. The PCA Component 2 values also show similarity among these samples, with only slight differences. The statistical data for the PCA have been attached (Supplementary File_Table S14).

**Figure 7:**
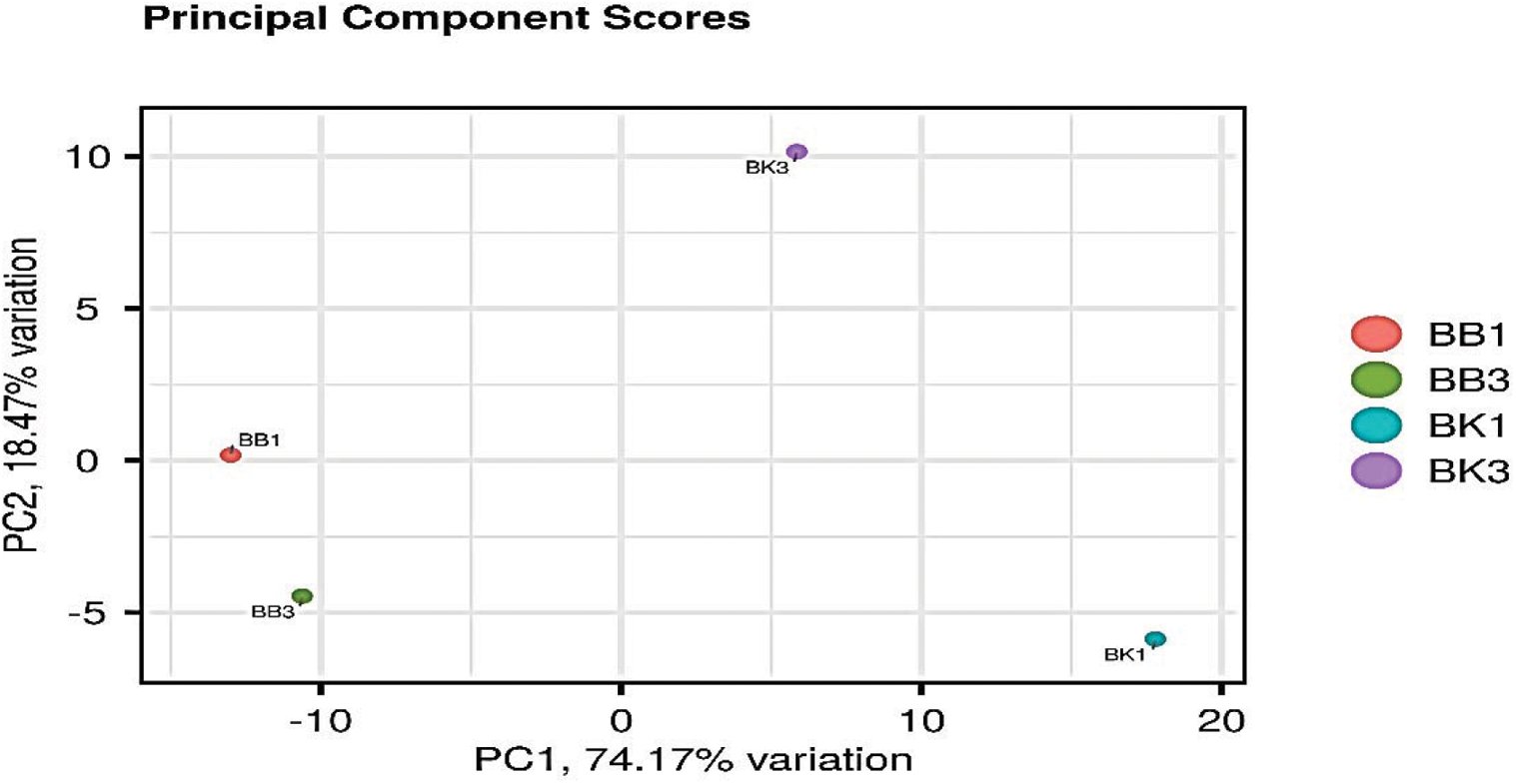
PCA analysis between BB and BK. PCA Component 1 & PCA Component 2 represents the similarity between BB1, BK1, BB3 & BK3.

### PEARSON CORRELATION BETWEEN BB and BK

On the basis of FPKM value, the Pearson correlation was measured between all the sheep group and across all the annotated genes. Pearson correlation is a statistical measure that evaluates the linear relationship between two continuous variables. In the context of sheep transcriptome, Pearson correlation can be used to identify genes that are co-expressed across different tissues or developmental stage.

**Figure 8:**
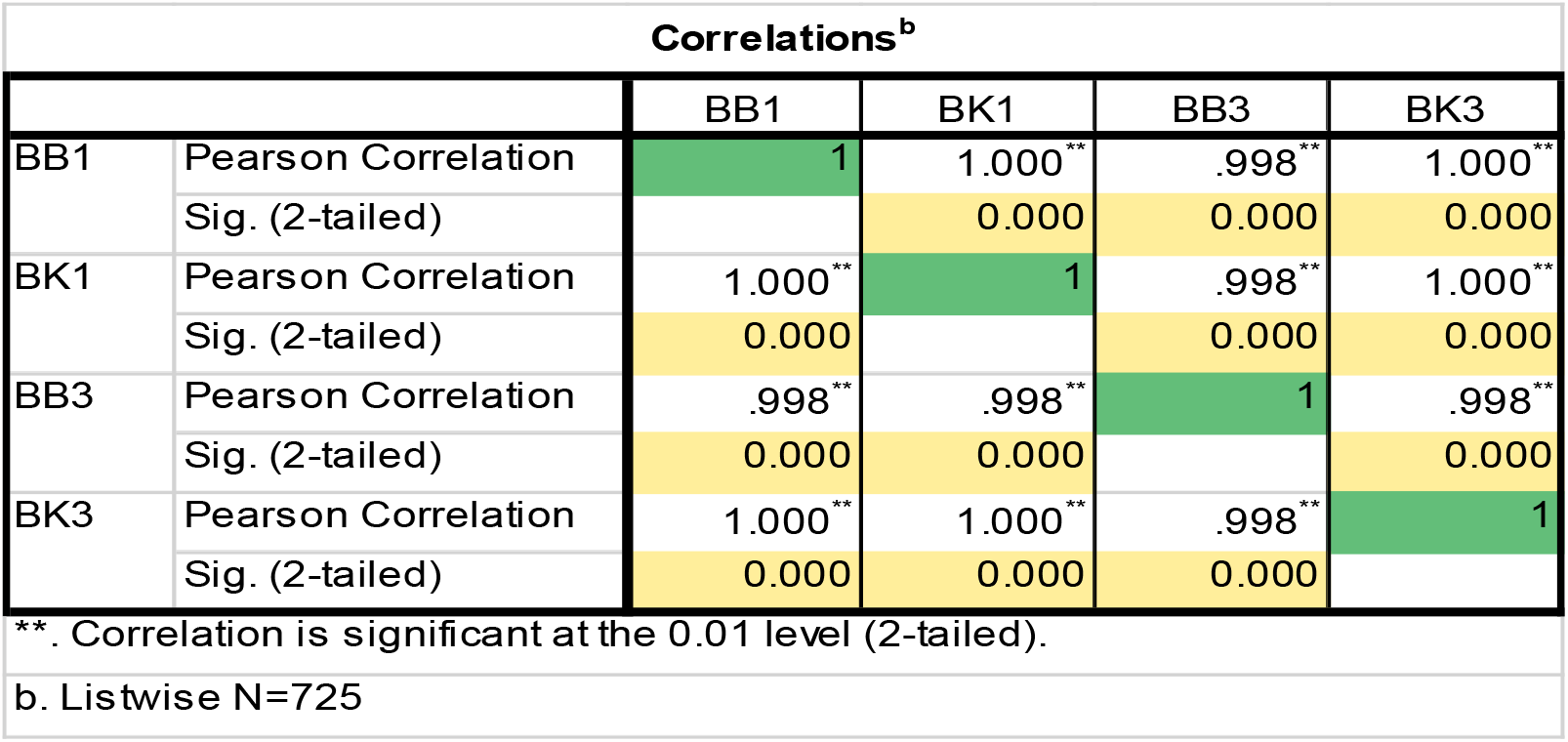
Pearson correlation between Kari and Balkhi based on their FPKM value.

### PROTEIN-PROTEIN INTERACTION (PPI) BETWEEN DEGS

The PPI were analyzed using the DEGs identifiers on high (700) confidence level. The edges indicate both functional and physical protein associations. These associations are meant to be specific and meaningful, i.e., proteins jointly contribute to a shared function; this does not necessarily mean they are physically binding to each other. The statistical data for the PPI have been attached (Supplementary File_Table S15-16).

**Figure 9:**
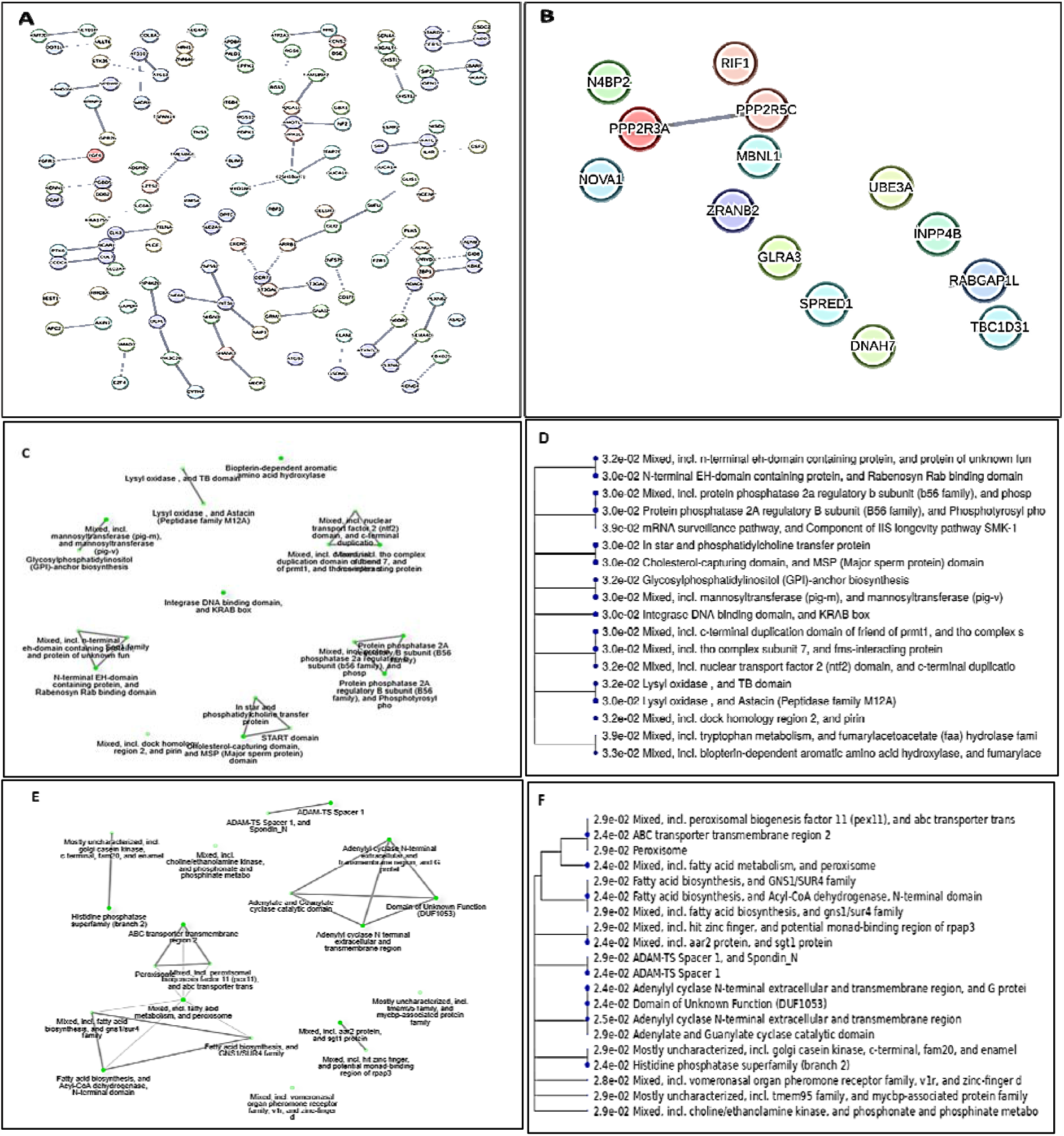
String function protein interaction network between the significantly (A) down-regulated genes and (B) up-regulated genes in BKvs.BB. (C) top 10 URGs and (D) tree. (E) Top 10 DRGs and (F) tree. Each node represents all the proteins produced by a single, protein-coding gene locus. Node colors represents interactions such as colored nodes: query proteins and first shell of interactors, white nodes: second shell of interactors. empty nodes: proteins of unknown 3D structure and filled nodes: a 3D structure is known or predicted.

## DISCUSSION

Reproductive traits contribute significantly in the world economy where lamb gestation is an important but complex process governed by a variety of factors. Kari is a unique sheep with a short gestation period [6]. The aim of the present study was to investigate the Kari sheep ovary transcriptome, including differentially expressed genes (DEGs), functional and Go analysis, which may be associated with gestational length. Several genes have been identified associated with reproductive traits in sheep [12].

In the ovary, transcriptome analysis revealed that genes expression of Kari and Balkhi was significantly different. Among the DEGs, we found GO terms: GO:0001228 associated with transcriptional activator activity, RNA polymerase II transcription regulatory region sequence-specific binding, and GO:0007416 linked with synapse assembly. The upregulated genes associated in these terms are *TCF12* controlling Transcriptional activator activity [13] and RNA polymerase, *CNOT4, RC3H1, XIAP* are thought to be associated with Ubiquitin-protein transferase activity [14] while *NFAT5, EPAS1, ZNF644, RBPJ, FOXP2* were shown to be associated with RNA polymerase II core promoter proximal region sequence-specific DNA binding [15]. A similar study on the transcriptomic of spermatogenesis in the testis of Hu sheep and Tibetan sheep was conducted by [16] to identify candidate genes and key pathways associated with fecundity in sheep. Their study identified several genes, including *COL1A1, COL1A2, COL3A1, SOX9, BCL2, HDC*, and GGT5, that were significantly enriched in terms and pathways that might affect the reproduction of sheep by regulating the migration of spermatogenic cells, apoptosis of spermatogenic cells, and secretion of sterol hormones via testicular interstitial cells. The study provides a theoretical basis for better understanding the molecular mechanisms of reproduction in sheep. These genes were highly expressed in BK and is reasonable to believe that they are linked with the gestation.

In addition, in our study, some up-regulated DEGs were significantly enriched in several metabolic KEGG pathways and organizational systems such as KEGG:3015 were associated with Translational activity, KEGG:532 linked with Metabolic pathways such as Glycosaminoglycan biosynthesis - chondroitin sulfate / dermatan sulfate. A similar study was conducted by [17] where the comparative transcriptomic analysis was applied to the liver and muscle of Dorper and Small-tailed Han sheep. Some DEGs, such as *TGFB1, TGFB3, FABP3, LPL*, and a number of GO terms may be associated with growth and fat deposition in Dorper.

Among the DEGs, we found three Go terms namely GO:0001228, GO:0004842 and GO:0000978 were associated with transcriptional activator activity”, “Ubiquitin-protein transferase activity” and “RNA polymerase binding activity. The down-regulated DEGs were significantly enriched in several metabolic KEGG pathways such as KEGG:4929 linked with endocrine system particularly GnRH secretion, KEGG:4550 associated with Cellular community – eukaryotes such as signaling pathways regulating pluripotency of stem cells. The downregulated genes included in these terms are *EEA1, CNOT4, FGD4, MBNL1, ZRANB2, REV3L, XIAP, ATP13A3, RPAP2, FOXP2, ADAMTS6*, these genes are associated with mRNA surveillance pathway [18]. A parallel study by [19] described a genome-wide comparison among ovine transcriptome derived from the M0, M3, M6, and M12 sheep testes by RNA-Sequencing. A number of key genes associated with reproduction were identified, which could be defined as candidate genes controlling testis development and spermatogenesis. These genes were highly expressed in BB and is reasonable to believe that they were linked with the gestation.

Also, in our study, the R value of the scatter plot indicates the correlation coefficient, which represents the strength and direction of the linear relationship between variables The R value between BB1 and BK1 is 0.06. A correlation coefficient of 0.06 is considered a weak correlation. Correlation coefficients typically range from -1 to 1, where -1 indicates a perfect negative correlation [20], 1 indicates a perfect positive correlation, and 0 suggests no correlation. In your case, a value of 0.06 indicates a very weak positive correlation [21], suggesting that there is a very slight, positive relationship between the gene expression patterns of the two sample groups. However, it’s not strong enough to draw robust conclusions, and the relationship may be influenced by various other factors. In summary, this PCA analysis suggests that the gene expression profiles of the four samples are quite similar, as they cluster closely along the first and second principal component axes, with slight distinctions between them.

In our analysis, we noticed an interesting pattern: Downregulated Genes (DRGs) showed higher expression levels in the control group compared to the experimental conditions. This suggests a reduction in gene expression in response to the experiment. On the other hand, Upregulated Genes (URGs) exhibited the opposite trend, with higher expression in the experimental group. This points to an increase in gene expression under specific experimental conditions. These findings offer valuable insights into how genes respond to different contexts, shedding light on the dynamic nature of gene regulation. The Protein-Protein Interaction (PPI) analysis offers valuable insights into the functional relationships among the identified DEGs. In the downregulated DEGs, *ABHD16B* and *NPBWR2* are the highly Co-expressed genes followed by *ADGRB2/GPR26*, and *AMOTL1/NF2*. While the upregulated highly Co-expressed DEGs were *DNAH7/TBC1D31, GLRA3I/NPP4B* and *MBNL1/NOVA1* respectively. A contrarily study by [22] suggested that The *MBNL1* and *NOVA1* SFs also tended to be negatively correlated with most Alternative Splicing (AS) events. In summary, identifying candidate genes and key pathways associated with gestation in sheep has important implications for genetic improvement, reproductive biology, sustainable agriculture, and animal welfare.

## CONCLUSION

Our study focused on the Kari sheep ovary transcriptome, focusing on differentially expressed genes (DEGs) associated with gestational length. DEGs were associated with GO terms such as “transcriptional activator activity” and “ubiquitin-protein transferase activity”. Notable genes were *TCF12, CNOT4, RC3H1*, and *XIAP*. Our correlation analysis showed a moderate positive correlation (R=0.06) between BB1 and BK1 gene expression patterns. DRGs in the control group showed higher expression, while URGs showed higher expression in the experimental group, shedding light on gene regulation dynamics Protein-protein interaction (PPI) analysis revealed a functional relationship between DEGs. This study enhances our understanding of pregnancy in sheep, indicating genetic development and reproductive biology.

## ETHICAL STATEMENT

The animal study was reviewed and approved by Animal Welfare and Ethics Committee of the Agriculture University Peshawar (Feb, 2021). Student Registration number is (2017-agr-u-37607).

## CONFLICT STATEMENT

The authors declare that the study is carried out in the absence of any commercial of financial relationships that could construct any potential conflict of interest.

### SIPPLEMENTARY MATERIAL

The supplementary data associated with this article have been attached as Supplementary_File.

